# Sound localization acuity of the common marmoset (*Callithrix jacchus*)

**DOI:** 10.1101/2022.08.13.503849

**Authors:** Evan D. Remington, Chenggang Chen, Xiaoqin Wang

## Abstract

The common marmoset (*Callithrix jacchus*) is a small arboreal New World primate which has emerged as a promising model in auditory neuroscience. One potentially useful application of this model system is in the study of the neural mechanism underlying spatial hearing in primate species, as the marmoset’s visually occluded natural habitat in the forest would make sound localization an essential behavior for survival. However, interpretation of neurophysiological data on sound localization requires an understanding of perceptual abilities, and the sound localization behavior of marmosets has not been well studied. The present experiment measured sound localization acuity using an operant conditioning procedure in which marmosets were trained to discriminate changes in sound location in the horizontal (azimuth) or vertical (elevation) dimension. Our results showed that the minimum audible angle (MAA) for horizontal discrimination was on average 15° for band-passed Gaussian noise and 13° for Random Spectral Shape (RSS) stimuli, whereas the MAA for vertical locations was at 17° and 22° for band-passed Gaussian noises containing more and less high frequency energy, respectively.

## 1. Introduction

The goal of the study of perception and cognition is to link brain activity with behavior. The common marmoset *(Callithrix jacchus)* has been a successful model for studying the auditory system in the past two decades (Wang 2018). The marmoset has been used to study the coding of pitch and complex spectral features in the auditory cortex (Bendor and Wang, 2005; Zhu et al., 2019; Zeng et al., 2021), temporal processing in the auditory cortex (Lu et al., 2001; Kajikawa et al., 2008; Zhou and Wang, 2010; Gao et al., 2016; Liu and Wang, 2022), thalamus (Bartlett and Wang, 2007), and inferior colliculus (Wang et al., 2022), spectral and intensity coding (Sadagopan and Wang, 2008; Song et al., 2022a), and sound source location (Remington and Wang, 2019; Zhou and Wang, 2014, 2012; Lui et al., 2015; Chen et al., 2022). Much is known about auditory cortex connectivity in marmosets (de la Mothe et al., 2012; Reser et al., 2009). Additionally, a number of studies have investigated auditory feedback mechanisms during vocalization (Eliades and Wang, 2008; Hage, 2020) and processing and control of conspecific communication in the prefrontal cortex (Roy et al., 2016; Jovanovic et al., 2022). As a model for hearing loss, marmosets have been used to study the representation of cochlear implant stimulation at the level of the auditory cortex (Johnson et al., 2012, 2016, 2017). Germline expression of a transgenic modification has been achieved in the marmoset (Park et al., 2016; Sasaki et al., 2009), broadening the marmoset’s potential as a model for neurologic diseases. There has also been an increasing interest in marmosets as a model for visual cognition (Mitchell et al., 2014; Davis et al., 2020; Song et al., 2022b).

Compared with the anatomy and physiology of the auditory system in marmosets, much less is known of their perceptual abilities. Our laboratory has developed an auditory operant conditioning task for the common marmoset to study auditory perception and for use in behaving neurophysiology (Remington et al., 2012). The task has been employed in the measurement of the audiogram (Osmanski and Wang, 2011), frequency discrimination thresholds (Osmanski et al., 2016), and harmonic resolvability and pitch perception (Osmanski et al., 2013; Song et al., 2016). Here we used this task to measure the spatial hearing acuity of the marmoset, specifically minimum audible angle (MAA).

As tropical arboreal species, marmosets need to navigate their environment using acoustic spatial cues. Spatial processing is therefore an important function performed by the marmoset’s auditory system, making the marmoset an ideal model species for further studies of spatial coding in the auditory cortex (Remington and Wang, 2019; Zhou and Wang, 2014, 2012). As in all mammals, sound location perception is determined by three cues: interaural time difference (ITD), interaural level difference (ILD), and spectral shape (Blauert, 1997). These cues are the result of the geometry of the head and ears: the distance between the ear canals determines ITD, the size and shape of the head (and to an extent the neck and shoulders) determines ILD, and the shape of the pinna (or outer ear) modulates the shape of the incoming sound spectrum by introducing resonances and notches in a spatially dependent manner. This spatially dependent acoustic filter is referred to as the head related transfer function (HRTF) (Blauert, 1997). This creates a dichotomy of perceptual computations for spatial perception: ITD and ILD are binaural cues and provide, at least in mammals, useful information about an object’s lateral position. A meta-analysis of measurements in multiple animal species finds that, at least for non-echolocating animals, there is a roughly linear relationship between head size and horizontal acuity (Brown and May, 2005), although there is also evidence that animals with highly focused binocular vision may also be localization specialists (Heffner and Heffner, 1992). Spectral cues, conversely, provide information related to front/back and up/down localization. It is appropriate, therefore, to measure acuity along these two axes independently. In this study, we measured the MAA of broad band sounds in azimuth and elevation. The results indicate that marmosets’ horizontal and vertical spatial acuity is approximately on par with other species of similar head size that have been previously tested.

## 2. Materials and methods

### 2.1 Subjects

Subjects were five common marmoset monkeys (2 male, 3 female) housed in individual cages in a colony at The Johns Hopkins University School of Medicine. Each animal was given free access to water; food was regulated during testing periods to keep animals at approximately 90% of free feeding weight. Subjects were tested once a day, five to seven days per week between the hours of 0900 and 1800. All experimental procedures were approved by the Institutional Animal Care and Use Committee of the Johns Hopkins University following National Institutes of Health guidelines.

### 2.2 Testing chamber and acoustic stimuli

Experiments were conducted in a double-walled sound-attenuated chamber (Industrial Acoustics, IAC, New York) with the internal walls, ceiling, and floor lined with ~3 inch acoustic absorption foam (Sonex, Illbruck). Acoustic stimuli were delivered from an array of 5-7 speakers (FT28D, Dome Tweeter, Fostex) mounted 1m from an animal’s head in a horizontal or vertical arc. Horizontal speakers (at 0-degree elevation) were placed in both frontal and rear locations. Vertical speakers were placed in the median plane in front of subjects (at 0-degree azimuth). Subjects sat in a wire mesh primate chair mounted onto a single stainless-steel bar such that the animals head was centered in the room (Remington et al., 2012). The primate chair was designed to minimize acoustic reflections near the pinna (**Fig. 1A**).

**Fig. 1.**
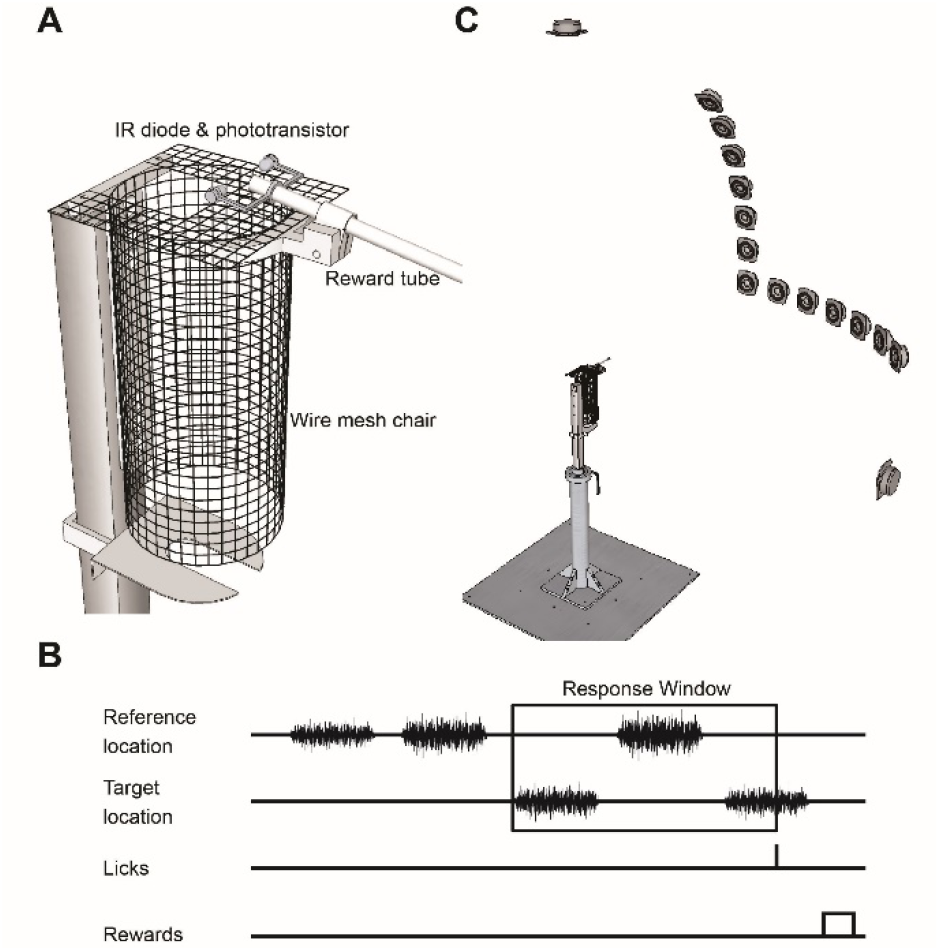
Marmoset chair and psychophysical testing procedure for measurement of minimum audible angle (MAA). **A.** Marmoset chair. Behavior sessions for MAA measurements were conducted while marmosets were seated in a steel wire chair designed to minimize acoustic reflections. Marmosets had to break a photobeam (photodiode & phototransistor) to register a response, which could be accomplished either by licking at the reward tube or moving the entire head forward. **B.** Location discrimination task trial structure. Marmosets listened to sounds (band-passed Gaussian noise or random spectral shape stimuli, see Methods) played from a reference location, and had to respond when a target location will began alternating with the reference location. If a response was registered within the preset number of alternations, a food reward was given. The next trial began after the animal finished consuming the reward (as measured via the photo beam). A response outside of a target interval resulted in a timeout. **C**. Speaker arrangement for MAA measurements. We measured localization discrimination thresholds in three conditions: frontal azimuth, rear azimuth, and vertically along the median plane. For rear azimuth discrimination, the chair was rotated 180°. Head position was monitored and with a closed-circuit camera system and custom image processing software (MATLAB). Marmosets were required to have heads facing forward in order for behavioral trials to proceed. Rear location is not pictured. The reference location was at 90-degree in azimuth (right lateral) or elevation (top) as shown.

Stimuli were generated in MATLAB (Mathworks, Natick, MA) at a sampling rate of 97.7 kHz using custom software. Digital signals were converted to analog (RX6, Tucker-Davis Technologies, Alachua, FL), then analog signals were attenuated (PA5, Tucker-Davis Technologies), power amplified (Crown Audio, Elkhart, IN), and played through a power multiplexer (PM2R, Tucker-Davis Technologies). Loudspeakers had a relatively flat frequency response curve (± 3-7 dB) and minimal spectral variation across speakers (< 7 dB re mean) across the range of frequencies of the stimuli used; all large (5-7 dB) spectral deviations occurred in narrow bandwidths near the upper limit of speakers’ frequency range (above 28 kHz), above the first spectral notch measured in marmoset head related transfer functions (Slee and Young, 2010).

Test stimuli included unfrozen (i.e., trial-unique) Gaussian noises that were band-pass filtered to contain energy between 2 and 32 kHz and Random Spectral Shape (RSS) stimuli (Barbour and Wang, 2003). RSS stimuli were constructed by summing pure-tones with pseudo-random levels centered around an average sound level. In this report all RSS stimuli were constructed to have energy between 2 and 32 kHz, with 30 tones per octave with levels varying independently in groups of three tones (i.e., 10 bins per octave). The standard deviation of the bin levels was 10 dB. Stimuli were 200 ms long with 10 ms cosine ramps. Sound level was roved ±10 dB to avoid the use of absolute level as a cue. The mean intensity was −35 dB relative to a maximum output intensity of 95 dB SPL at 0 dB attenuation for a 4 kHz tone.

### 2.3 Psychophysical testing procedure

Marmosets were trained to sit in the custom-designed primate chair and perform a Go/No-Go spatial location discrimination task (Brown et al., 1980) using the method of constant stimuli (Gescheider, 1985). Details of behavioral training and behavioral apparatus have been described previously (Osmanski and Wang, 2011; Remington et al., 2012). Behavioral responses involved breaking a photo beam attached to a feeding tube, either by moving the entire head slightly forward into the beam or with licks at the tube (**Fig. 1A**). Each behavior session was composed of a preset number of trials (typically 80-100), where each trial was composed of a variable duration ‘inter-target interval’ and a fixed duration ‘response interval.’ Inter-target interval duration was randomized between approximately 3 and 10 seconds. The response interval was dependent on the number and duration of targets and was typically 5 seconds in length. During an inter-target interval, a series of reference sounds were played from a fixed location with a 700 ms inter-stimulus interval. The duration of reference and target sounds were both 200 ms. Behavioral responses during this time result in a mild punishment (such as a timeout or a puff of air at the base of the tail) and a restarting of the trial. If an animal withheld response during the inter-target interval, the sound location began alternating between one of several target locations and the reference location during the response interval. A trial ended when the response interval expired or a response was detected during the response interval. Behavioral responses during this time were reinforced with a small food reward (**Fig. 1B**). Approximately 30% of trials were “catch trials,” which were identical in length to target trials in their timing and structure but in which no targets are delivered (i.e., sounds continued to be delivered from the reference speaker). Thus, during a catch trial the response interval is indistinguishable from the inter-target interval from the animal’s perspective. A response during a catch trial response interval is referred to as a false alarm (or false positive).

Head position was tracked using an infrared camera and a custom algorithm written for MATLAB using the image processing toolbox. Prior to each stimulus delivery, the algorithm checked to see if the eyes could be located in an experimenter-defined region. Therefore, animals had to face forward in order for behavioral trials to proceed. If an animal closed its eyes, the algorithm would fail to find the eyes and the program would pause. In addition to assuring that animals maintained open eyes, a specific head position can be considered as indication that an animal is engaged in the task.

To quantify behavior performance for the measurement of MAA in marmosets, we constructed psychometric functions over a fixed set of distances (i.e., target locations) from the reference location (**Fig. 1C**). The target locations were chosen for each MAA measurement based on animals’ performance early in testing such that hit rates were sufficiently high to motivate animals to perform the task. In some cases, the hardest targets were omitted for certain animals. Thresholds were determined to be the point at which the interpolated psychometric function equaled 50%. To adjust for false positives, a corrected hit rate was used:

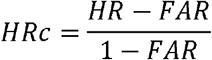

where {HR} is the raw hit rate, and {FAR} is the false alarm (or false positive) rate (Gescheider, 1985). Final thresholds were calculated from psychometric functions averaged over 3 of 4 consecutive sessions during which thresholds were no higher than the mean plus 5° and during which time thresholds did not appear to be systematically decreasing. Sessions with a false alarm rate greater than 30% were not included in threshold calculations. Not all animals were tested in each condition.

## 3. Results

### 3.1 Horizontal acuity

For all noise stimuli, noise tokens were generated uniquely for every trial. The ability of marmosets to discriminate sound source azimuth of band-pass filtered Gaussian noise stimuli is illustrated in **Fig. 2A**. The five animals tested showed generally good agreement in thresholds, with a mean threshold of 15° and standard deviation of 4°. Most of the variation among the subjects occurred at 15° separation; most could reliably discriminate 22.5° separation, yet none could do so for 7.5° separation.

**Fig. 2.**
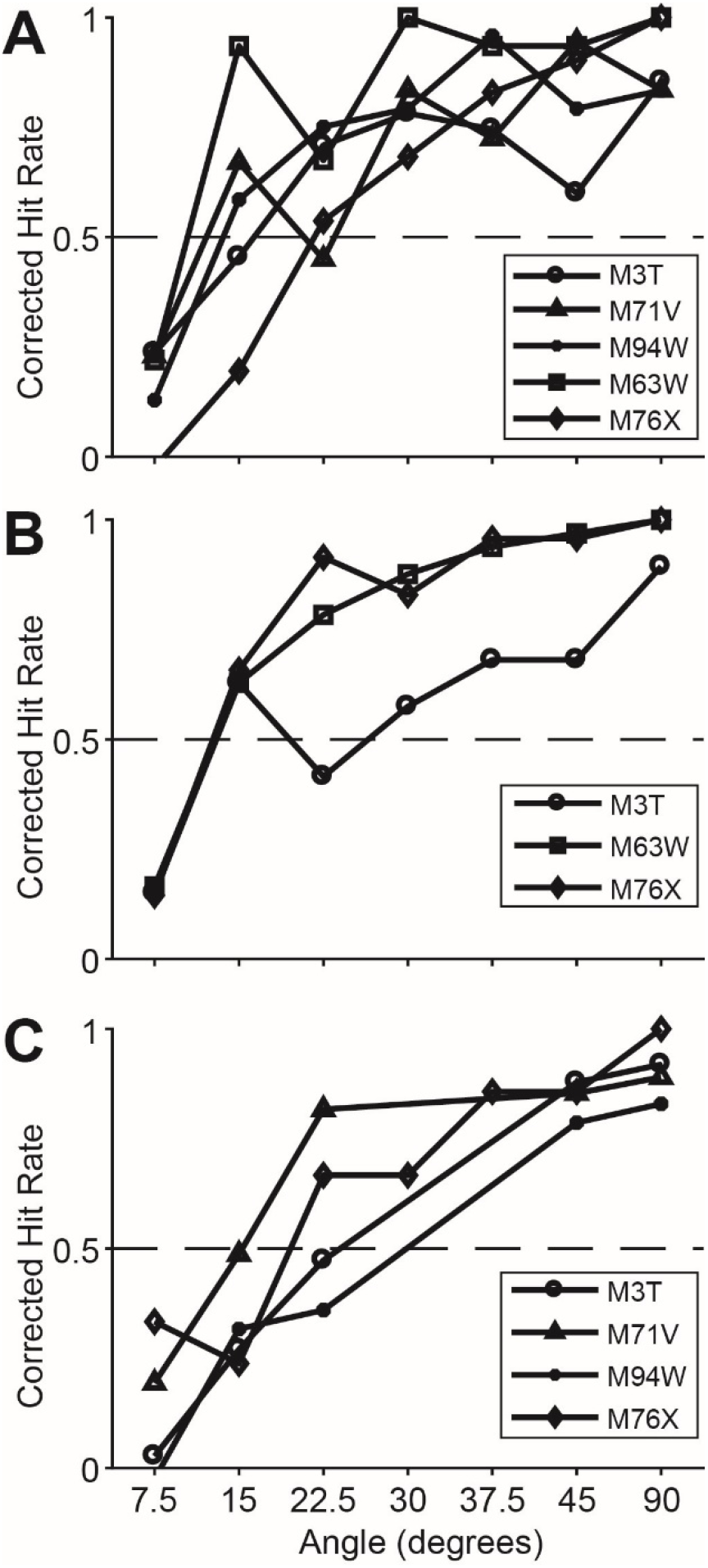
Psychometric functions for horizontal location discrimination for different stimuli. In all plots, data points represent averaged corrected hit rate (see Methods) to different angles between the reference location and target locations. Different marmosets are represented by different marker shapes, as shown in the legend (right). **A.** Gaussian noise, band passed between 2 and 32 kHz. **B.** RSS stimuli, containing energy between 2 and 32 kHz. For RSS, the frequency profile varied between stimuli, making it difficult to discriminate stimuli on the basis of monaural spectral cues. **C**. Psychometric functions for rear locations, stimuli were 2-32 kHz Gaussian noise.

Although sound localization in the horizontal plane relies primarily on binaural cues, HRTF shape has also been shown to change with azimuth in several species (Wightman & Kistler 1989; Rice et al. 1992). However, these changes are mostly confined to higher frequency cues in marmosets (Slee and Young, 2010), and cats have been shown to not depend heavily on specific frequency cues in a horizontal discrimination task (Huang and May, 1996a). Still, there is a possibility that in addition to binaural cues, monaural spectral cues could be used to perform a horizontal discrimination task (Butler, 1986; Van Wanrooij and Van Opstal, 2004). To reduce the possibility that marmosets could be using spectral cues for azimuth discrimination, we tested three marmosets’ ability to discriminate the location of random spectral shape (RSS) stimuli which varied spectrally on each stimulus presentation and made it difficult to compare successive stimuli on the basis of frequency spectrum. We hypothesized that if subjects were relying on spectral cues to perform azimuth discrimination, thresholds should be higher when discriminating RSS stimuli. Psychometric functions for RSS stimuli are shown in **Fig. 2B**. Thresholds were as low (mean 13°), if not lower, as in the Gaussian noise condition, suggesting that the marmosets tested used binaural cues to discriminate horizontal locations.

Data from previous studies suggest that localization is less accurate at rear locations compared with frontal locations (Oldfield and Parker, 1984; Recanzone and Beckerman, 2004). However, because marmosets are a tropical arboreal species living mostly in a visually occluded natural environment with dense forest, we thought it would be important to test whether they possessed relatively heightened localization abilities outside the visual field. We tested four marmosets’ ability to discriminate azimuth at rear locations using Gaussian noise stimuli. These results are shown in **Fig. 2C**. Animals were not tested as extensively in this condition as the front location, however, thresholds (mean 22°) appeared to be generally elevated when compared with the front locations, indicating that, at least without additional training, rear horizontal acuity in marmosets is worse than frontal horizontal acuity.

### 3.2 Vertical acuity

While horizontal spatial information is contained in the binaural differences between the two ears, additional information is required to compute sound source location in a 3-dimensional space. To do this, the auditory system makes use of spectral cues generated primarily by the pinna. These cues in marmosets are located at frequencies higher than 12 kHz (Slee and Young, 2010). We measured vertical location acuity first using the same Gaussian noise stimuli used to test horizontal acuity. Psychometric functions for three animals are shown in **Fig. 3A**. The average threshold was 17°.

**Fig. 3.**
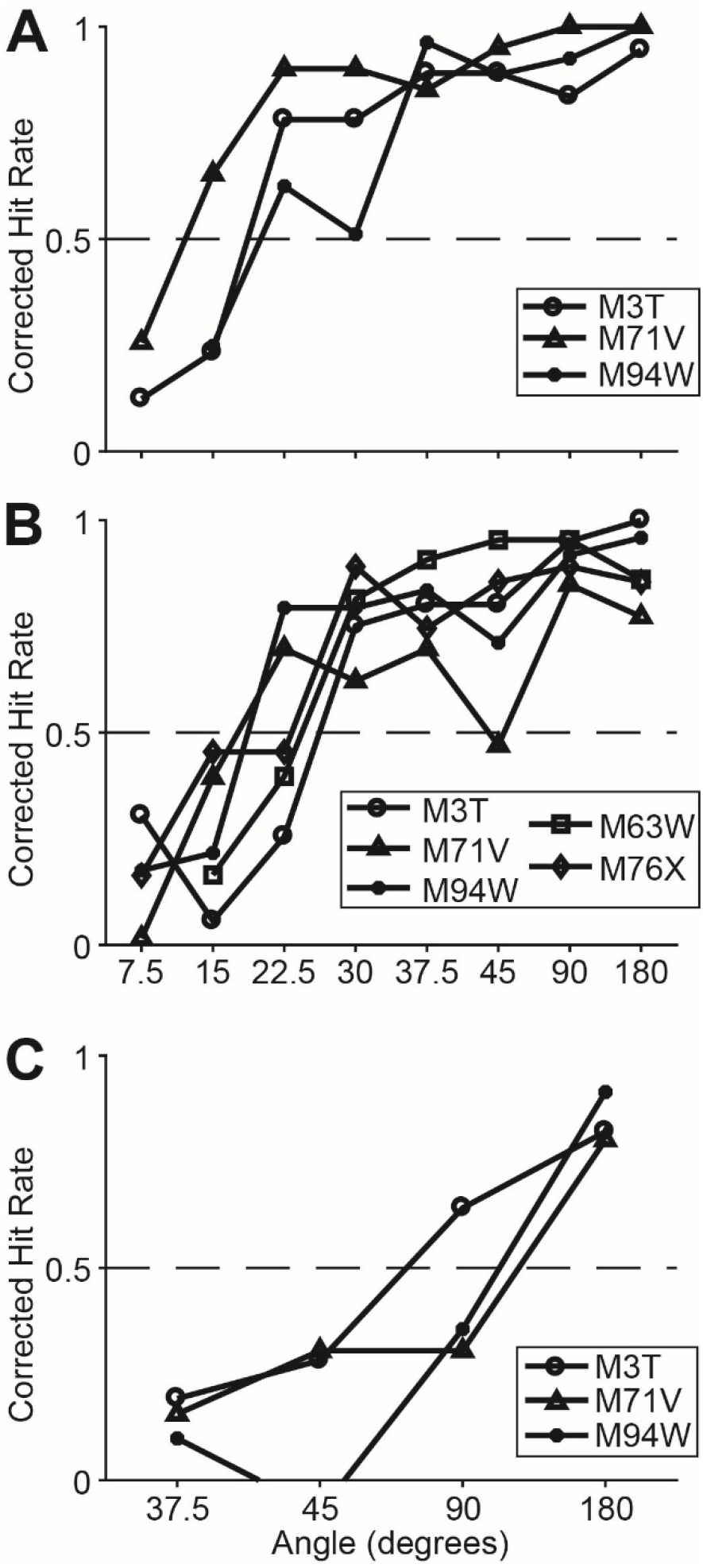
Psychometric functions for vertical locations discrimination for different stimuli. Format same as **Fig. 2**. **A.** Gaussian noise, band pass filtered between 2 and 32 kHz. **B**. Gaussian noise, filtered between 4 and 26 kHz to exclude acoustic information above the first spectral notch measured previously in marmoset HRTFs (see Results). Thresholds were not substantially higher than for 2-32 kHz Gaussian noise, indicating that vertical discrimination acuity was not dependent on very high frequency cues. **C**. Gaussian noise, filtered between 4 and 12 kHz to exclude acoustic information in the range of the first spectral notch. Localization acuity was much lower for this stimulus. M94W had a negative corrected hit rate at 45 degrees. This was because the hit rate (HR) was lower than the false alarm rate (FAR) at 45 degrees. Notice that only sessions with a false alarm rate lower than 30% were included in threshold calculations.

The HRTF of mammals contains several components which could be useful for directional hearing. These include a “mid-frequency” component, in which there exists a prominent spectral notch (sometimes called the “first notch”) that varies in frequency in a somewhat predictable manner with elevation, and a “high frequency” component, which inhabits the upper range of audibility and contains cues that can vary in a less predictable manner (Rice et al., 1992; Slee and Young, 2010). In marmosets, this first notch was shown to vary between 12 and 24 kHz (Slee and Young, 2010). To reduce the possibility that marmosets were using high frequency cues, vertical discrimination acuity was measured in several animals using Gaussian noises filtered between 4 and 26 kHz. Psychometric functions for five marmosets discriminating elevation using the mid-frequency Gaussian noise stimuli (4-26 kHz) are shown in **Fig. 3B**. The average threshold was 22°, slightly higher than when the stimuli included higher frequency information up to 32 kHz, but the difference was not significant (rank-sum test, p = 0.29). Three animals were tested using both types of stimuli. Two of the animals displayed higher thresholds when tested by the mid-frequency Gaussian noise stimuli (M3T: 19° vs. 26°; M71V: 12° vs. 18°), whereas one had slightly lower thresholds (M94W: 20° vs. 19°).

We additionally tested three animals using sounds which were filtered to only include stimulus information below the location of the first spectral notch (**Fig. 3C**) (Gaussian noise, 4-12 kHz). Animals tested with this stimulus exhibited extreme difficulty in discriminating vertical location. One animal had a threshold of 72°, while other two animals could not discriminate any of the locations in the frontal hemifield (112° and 124°). All three animals, however, could reliably discriminate front and rear locations (i.e., threshold < 180°).

## 4. Discussion

### 4.1 Horizontal sound location discrimination

The average threshold for horizontal localization of 15° (13° for RSS stimuli) measured in marmosets is higher than many other mammals. Several species have been shown to discriminate sound locations separated by less than 10° in azimuth, including cats (Heffner and Heffner, 1988), macaques (Brown et al., 1980), and opossums (Ravizza et al., 1972). Despite significant training, 10° seemed to be unachievable for azimuth discrimination by marmosets in the current study. It is unlikely that this limit was due to a lack of motivation, as false positive rates were not particularly low; all animals had false positive rates higher than threshold on several occasions. Another possibility is the use of sounds filtered above 2 kHz, which is above the frequency region where ITD cues are most useful. In humans, the upper limit for ITD cues has been measured at 1.3 kHz (Klumpp and Eady, 1956). We believe this is also an unlikely limiting factor, as cats and macaques have both been shown to exhibit high horizontal acuity using test stimuli of comparable spectra (Brown et al., 1980; Huang and May, 1996a). Also, there is some evidence that several small mammals (tree shrew, rat, gerbil) can use ITD cues at frequencies significantly higher than 2 kHz (Heffner and Masterton, 1980; Masterton et al., 1975). This is possibly to compensate for the lack of strong ILD cues in the middle frequency region due to small head size. In fact, fine structure phase locking in the auditory nerve has been observed at frequencies up to 4 kHz in both cats and squirrel monkeys (Johnson, 1980; Rose and Brugge, 1967). In marmosets, ILD cues are relatively small below 5 kHz (Slee and Young, 2010).

We believe a better explanation for the higher horizontal threshold observed here in marmosets compared to those in cats and macaques therefore is the relatively small head size of marmosets. As binaural cues are known to be dependent on the physical distance between the two ears (for ITD) and the size of the head (for ILD), it is perhaps not surprising that marmosets do not appear to be expert localizers. A meta-analysis of data from multiple species shows a relatively good correlation between head size (as defined by maximum interaural time difference) and horizontal discrimination threshold (**Fig. 4A**) (Brown and May, 2005). Placing the marmoset into this dataset shows that the performance measured in the present study is almost exactly what would be predicted based on head size.

**Fig. 4.**
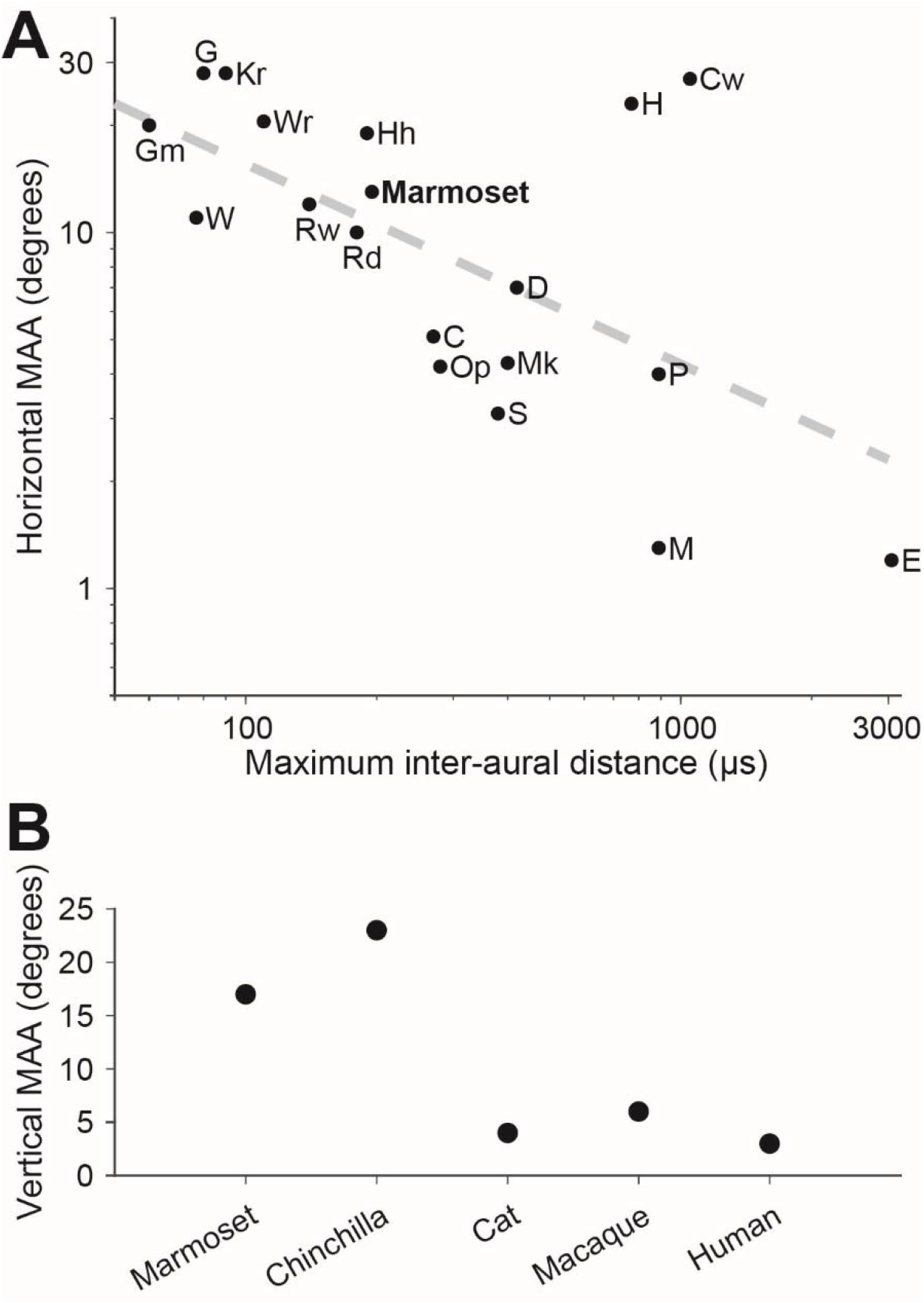
Comparative sound localization acuity. **A**. Sound localization thresholds for broad-band stimuli as a function of head size in 18 mammals, plus the present value for marmosets. Marmosets are roughly in line with the trend gathered from previous studies (diagonal line, r = −0.59). Gm, grasshopper mouse (Onychomys leucogaster); W, least weasel (Mustela nivalis); G, gerbil (Meriones unguiculatus); Kr, kangaroo rat (Dipodomys merriami); Rw, wild Norway rat (Rattus norvegicus); Rd, domestic Norway rat and Wistar albino rat (R. norvegicus) Wr, wood rat (Neotoma floridiana); Hh, hedgehog (Paraechinus hypomelas); C, cat (Felis catus); Op, opossum (Didelphis virginiana); S, harbor seal (Phoca vitulina); Mk, rhesus and pig-tailed macaque monkeys (Macaca mulatta) and (M. nemestrina); D, dog (Canis canis); H, horse (Equus caballus); M, human (Homo sapiens); P, domestic pig (Sus scrofa); Cw, cattle (Bos taurus); E, elephant (Elephas maximus). Figure reproduced and modified, with permission, from Brown and May (2005). **B**. Vertical sound localization discrimination thresholds in several mammals.

Although sound localization in the horizontal plane relies primarily on binaural cues, HRTF shape has also been observed to change with azimuth in several species (Rice et al., 1992; Wightman and Kistler, 1989), and monaural spectral cues can be used to perform a horizontal discrimination task (Butler, 1986; Van Wanrooij and Van Opstal, 2004). As this task measured discrimination ability rather than absolute localization accuracy, the usefulness of these spectral cues could be increased. Our results showed that RSS thresholds were equally low or lower than those measured in the Gaussian noise condition (**Fig. 2B)**, suggesting that marmosets used primarily or exclusively binaural cues to discriminate horizontal locations.

Although binaural cues are often weaker at rear locations, and data suggest that localization is less accurate at rear locations compared with frontal locations (Oldfield and Parker, 1984; Recanzone and Beckerman, 2004), an animal suited to an arboreal environment might possess heightened sensitivity to space outside the visual field. The observations that rear acuity was lower when compared with front locations indicated that this may not be the case for marmosets (**Fig. 2C)**.

### 4.2 Vertical sound localization

A comparison of marmoset vertical acuity with other tested animals is shown in **Fig. 4B**. Our results showed that marmosets’ thresholds for vertical localization were higher than those for horizontal localization. This finding is consistent with several species previously tested, such as the chinchilla (Heffner et al., 1995) and the opossum (Ravizza et al., 1972). Vertical acuity in several species, such as cats (Martin and Webster, 1987) and macaques (Brown et al., 1982) has been shown to be much better (roughly equal to horizontal acuity) than in marmosets. The exceptional vertical discrimination in these species, however, is degraded by the removal of high frequency energy (Brown et al., 1982; Huang and May, 1996a). The HRTF of mammals includes a “mid-frequency” component, in which there exists a prominent spectral notch that varies in frequency in a somewhat predictable manner with elevation, and a “high frequency” component, which inhabits the upper range of audibility and contains cues that can vary in a less predictable manner (Rice et al., 1992; Slee and Young, 2010). In cats, accurate absolute localization (i.e., localization identification by head orientation) is dependent on mid-frequency cues, degrading when only high frequency cues are available (Huang and May, 1996b), thus vertical discrimination thresholds using stimuli with high frequency energy may overestimate the actual localization ability of listeners. In cats discriminating “mid-frequency” sounds (5-18 kHz), vertical localization thresholds were shown to be higher than azimuth thresholds (Huang and May, 1996a).

In marmosets, the first notch in HRTF varies between 12 and 24 kHz (Slee and Young, 2010). To sort out contributions by spectral cues in different frequency ranges, we measured the vertical acuity using three sets of Gaussian noise stimuli (**Fig. 3**). The initial test stimulus, with a frequency range of 2-32 kHz, included energy in marmosets’ “high frequency” region. To test acuity without access to information at high frequency cues outside the range of the first notch, vertical discrimination acuity was tested in several animals using Gaussian noise filtered between 4 and 26 kHz (mid-frequency). The average threshold of 22° was slightly higher than when the stimulus included higher frequency information. Finally, to test whether marmosets were in fact using energy in the first notch region and not using low frequency cues (lower than 12 kHz), we measured the vertical acuity in two marmosets using a 4-12 kHz Gaussian noise. Performance was poor using this stimulus, indicating that marmosets did not use low frequency cues to perform vertical location discrimination.

## Abbreviations

MAA: minimum audible angle;
HRTF: head-related transfer function

## CRediT authorship contribution statement

**Evan D. Remington:** Conceptualization, Methodology, Software, Investigation, Data curation, Visualization, Writing – original draft. **Chenggang Chen:** Visualization, Writing – review & editing. **Xiaoqin Wang:** Conceptualization, Funding acquisition, Resources, Writing – review & editing, Supervision.

## Acknowledgements

We thank J. Estes and N. Sotuyo for assistance with animal care, C. Baldwin, J. Green, M. Gu, and T. Saavedra for generous assistance in performing some of the experiments, and M. Osmanski for his comments on the manuscript.

